# DrugHIVE: Target-specific spatial drug design and optimization with a hierarchical generative model

**DOI:** 10.1101/2023.12.22.573155

**Authors:** Jesse A. Weller, Remo Rohs

**Affiliations:** Department of Quantitative and Computational Biology, University of Southern California, Los Angeles, CA 90089, USA; Department of Physics and Astronomy, University of Southern California, Los Angeles, CA 90089, USA; Department of Chemistry, University of Southern California, Los Angeles, CA 90089, USA; Thomas Lord Department of Computer Science, University of Southern California, Los Angeles, CA 90089, USA

## Abstract

Rapid advancement in the computational methods of structure-based drug design has led to their widespread adoption as key tools in the early drug development process. Recently, the remarkable growth of available crystal structure data and libraries of commercially available or readily synthesizable molecules have unlocked previously inaccessible regions of chemical space for drug development. Paired with improvements in virtual ligand screening methods, these expanded libraries are having a significant impact on the success of early drug design efforts. However, screening-based methods are limited in their scalability due to computational limits and the sheer scale of drug-like space. An approach within the quickly evolving field of artificial intelligence (AI), deep generative modeling, is extending the reach of molecular design beyond classical methods by learning the fundamental intra- and inter-molecular relationships in drug-target systems from existing data. In this work we introduce DrugHIVE, a deep hierarchical structure-based generative model that enables fine-grained control over molecular generation. Our model outperforms state of the art autoregressive and diffusion-based methods on common benchmarks and in speed of generation. Here, we demonstrate DrugHIVE’s capacity to accelerate a wide range of common drug design tasks such as de novo generation, molecular optimization, scaffold hopping, linker design, and high throughput pattern replacement. Our method is highly scalable and can be applied to high confidence AlphaFold predicted receptors, extending our ability to generate high quality drug-like molecules to a majority of the unsolved human proteome.

## INTRODUCTION

Success in drug development depends largely on the quality of initial candidate molecules identified during the early drug discovery process^1,2^. Historically, a major obstacle to finding better candidates has been the limited size of drug-like molecular libraries which until recently have been limited to a few million compounds. In recent years, the number of commercially available or readily synthesizable drug-like chemicals have grown into the billions of compounds^3–5^, though this remains a miniscule fraction of the total drug-like chemical space—estimated to be as large as 10^60^ compounds^6^. Structure based virtual ligand screening (VLS) approaches have also been limited until recently to well below 10^7^ compounds. With the advent of virtually enumerated libraries and fragment based virtual screening techniques^7,8^, this capacity has increased into the billions and will possibly extend to the tera-scale in the near future^9^. These techniques are having impact in accelerating and lowering the cost of early drug discovery by drastically growing the initial candidate pool and mitigating the need for expensive large-scale high throughput screening (HTS). However, they still rely on the explicit enumeration of chemical libraries and therefore will not be readily scalable to significant fractions of the almost infinite drug-like space^10^. Other critical steps in the early drug development process are limited due to their manual nature, such as the systematic modification of candidates during hit-to-lead optimization. Although progress has been made in leveraging computational tools applying structure-based drug design (SBDD)^11^, these costly and time-consuming tasks still rely significantly on, and are therefore limited by, the expertise and intuition of human intelligence^2^. Overcoming such limitations and expanding the reach of computational drug design to greater scales of chemical space will require new approaches.

Recent advances in artificial intelligence and generative modeling, which have yielded impressive results on a number of previously intractable problems in biology^12,13^, provide a promising new approach. Rather than further optimizing the exhaustive search of chemical space, deep generative models are capable of learning distributions over molecular data and have already been successfully applied to large drug-like chemical datasets for de novo molecular generation^14–18^.Lately, efforts have turned toward modeling drug-target interactions by training models on datasets of ligand-bound receptor crystal structures^19–22^. By learning a joint probability distribution of ligands bound to their receptors, these models can generate new molecules conditioned on the receptor, effectively narrowing the regions of chemical space in which to search for new drugs. As the field progresses, generative models are being designed with new capabilities such as scaffold hopping, fragment growing, linker design, and molecular property optimization, all to better meet the challenges of early drug design. Perhaps the most important benefit of the generative modeling approach is the capacity to generalize the drug-target interactions learned from existing data to new receptor structures. This is in stark contrast to the classical VLS and HTS approaches in which the significant time and cost of screening large molecular libraries must be repeated with each new target.

In this work, we propose DrugHIVE (Figure 1), a deep structure-based <u>Drug</u> generating <u>HI</u>erarchical <u>V</u>ariational auto<u>E</u>ncoder. Unlike previous encoder-decoder models, our approach employs a hierarchical prior structure which represents more naturally the distribution of molecular space. The use of a hierarchical prior leads to the encoding of input molecular structures at varying spatial scales and enables the generation of new molecules with a high level of spatial control—a necessity for many drug design tasks. We show that DrugHIVE outcompetes state of the art autoregressive and diffusion-based approaches in generating molecules with predicted high binding affinity and drug-likeness scores. Also, since it generates molecules in a rapid single-shot fashion, our model is significantly faster than these other approaches which require multi-step inference and tend to be notoriously slow^23^.

**Figure 1:**
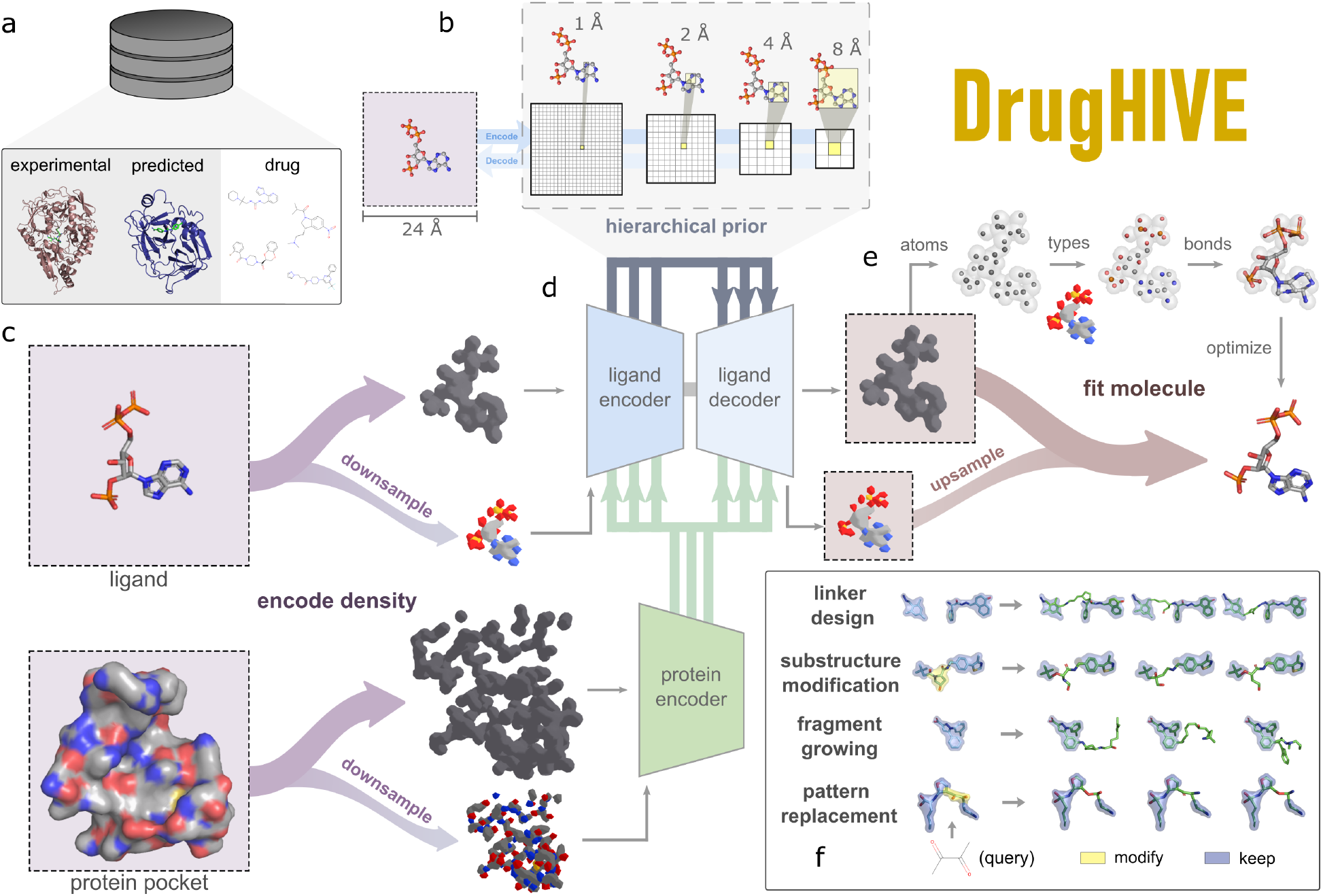
Overview of DrugHIVE. **(a)** Input data to the model can come from experimental structures, predicted structures, or drug-like molecule libraries. The model is trained on data from experimental structures and drug-like molecules, and generation can be performed using any of the three data types. **(b)** The prior of DrugHIVE has a hierarchical structure, with varying spatial scales being represented. **(c)** Molecular information is the input of the model in the form of multi-channel atomic density grids, with ligand grids input into the ligand encoder and receptor grids input into the protein encoder. Atomic occupancy (dark grey density) is represented at full grid resolution whereas atom type channels (colored density) are downsampled by a factor of 2 before being input into the network. **(d)** The DrugHIVE model is comprised of a ligand encoder, a protein encoder, and ligand decoder. Information from the protein encoder is passed into both the ligand encoder and decoder. **(e)** The model outputs atomic occupancy (full resolution) and atom type grids (half resolution) which are then fit with a molecular structure. First, atoms are fit to the atomic occupancy density and atomic types are assigned using the upsampled atom type density, then a bond fitting algorithm is applied to connect the atoms. Finally atomic positions and bond lengths are relaxed using force field optimization. **(f)** DrugHIVE can automate many common drug design tasks such as linker design, substructure modification (scaffold hopping), fragment growing, and pattern replacement.

Owing to its hierarchical structure and readily accessible encoding space, DrugHIVE can generate molecules in myriad ways and automate a wide range of common tasks in drug design. We show an impressive ability to optimize the drug-like properties, binding affinity, and selectivity of molecules through evolutionary latent space search. We apply our model to the automated replacement of Pan-Assay Interference Compounds (PAINS) patterns, the linking of fragments from a fragment screening experiment, substructure optimization (scaffold hopping), and the growing of molecular structures (fragment growing). Finally, we demonstrate that our model can successfully generate and optimize ligands with improved predicted binding affinity for receptors using AlphaFold2 (AF) predicted structures, extending the limits of generative molecular design beyond the set of currently available crystal structures.

## RESULTS

### De novo drug generation with prior sampling

DrugHIVE excels in generating new molecules with drug-like properties and a strong predicted binding affinity for their target receptors, outperforming diffusion and graph-based networks in the average predicted binding affinity and drug-likeness of generated molecules. We test the ability of DrugHIVE to generate new candidate ligands for a test set of 100 diverse receptors and compare the results against competitors and a random sample of molecules from the ZINC drug-like subset (Figure S2). For each receptor, we generate molecules by sampling randomly from the latent prior and then compute the estimated binding affinity and commonly reported properties of each molecule. Our model performs well against its competitors on common benchmark metrics. The molecules generated by our model have better predicted binding affinities and drug-likeness scores, on average, than competitive state of the art models. Approximately 10.3% of the DrugHIVE generated ligands have a higher predicted affinity than the reference ligand. This is about double the value (4.6%) of a random sample of the ZINC dataset, which means that screening efficiency is significantly higher for DrugHIVE generated molecules.

Though it has become customary in the literature to report these metrics as a benchmark of generative performance, there have been some common oversights regarding how they are calculated and interpreted. For instance, most papers in the field neglect to correct for the strong correlation of Vina score, the most commonly used metric for binding affinity, with molecule size. This means that the best models reported in the recent literature may simply be the ones that generate the largest molecules. This is particularly important to correct for given that many models, such as point-cloud and graph-based approaches, require the size each generated molecule to be specified. It is also common for studies to report the average diversity of generated molecules as a metric to be uniformly maximized, where the best model is considered to be the one that generates the highest molecular diversity. This logic may mostly hold for the generation of raw molecules, but it breaks down for ligands conditioned on specific target receptors. For example, given the specificity of drug-target interactions, we would not expect a good model to generate molecular diversity significantly higher than a random set of drug-like molecules from ZINC. Therefore, within a reasonable range of average diversity values, it will be unclear which is best.

To further demonstrate, we generate 1,000 molecules for the Rho-associated protein kinase 1 (ROCK1) receptor (PDB ID 3TWJ), an important target with a documented role in cancer migration, invasion, and metastasis^24^. Approximately 12% of the generated ligands have a higher predicted binding affinity than the crystal ligand (RKI-1447), a type I kinase inhibitor with reported IC50=14.5 nM. In Figure S1, we compare the properties of the reference ligands in the PDBbind^25^ dataset and a random sample of the molecules from the ZINC^5^ dataset with those of the generated ligands from DrugHIVE. The properties of the generated molecules fall well within the distributions of the training datasets, which is an indicator that the model effectively learns the training distributions. We also observe a significant distribution shift between the molecules generated with and without protein receptors, which indicates that the model is indeed conditioning on receptor information during generation.

### Multi-objective optimization of drug properties via evolutionary latent search

DrugHIVE can generate new ligands with a high degree of control over similarity to a reference molecule using prior-posterior sampling (Figure 2f). Each ligand-receptor complex has a unique representation in the encoded latent space, so if we start with a given reference ligand latent representation (posterior) and interpolate toward a randomly sampled point (prior), the resulting molecules will have a higher average diversity and lower molecular similarity to the reference as we interpolate toward the prior sampled point (Figure 2d). Similarly, the distribution of binding affinities shifts away from that of the reference ligand and significantly widens (Figure 2e). Taking advantage of this high level of control over variation, we implement an evolutionary algorithm to search the latent space and optimize the properties of a reference molecule. In Figure 2a-c we show the optimization process for the ligand Lisinopril bound to the human angiotensin-converting enzyme (ACE) receptor (PDB ID 1O86).

**Figure 2:**
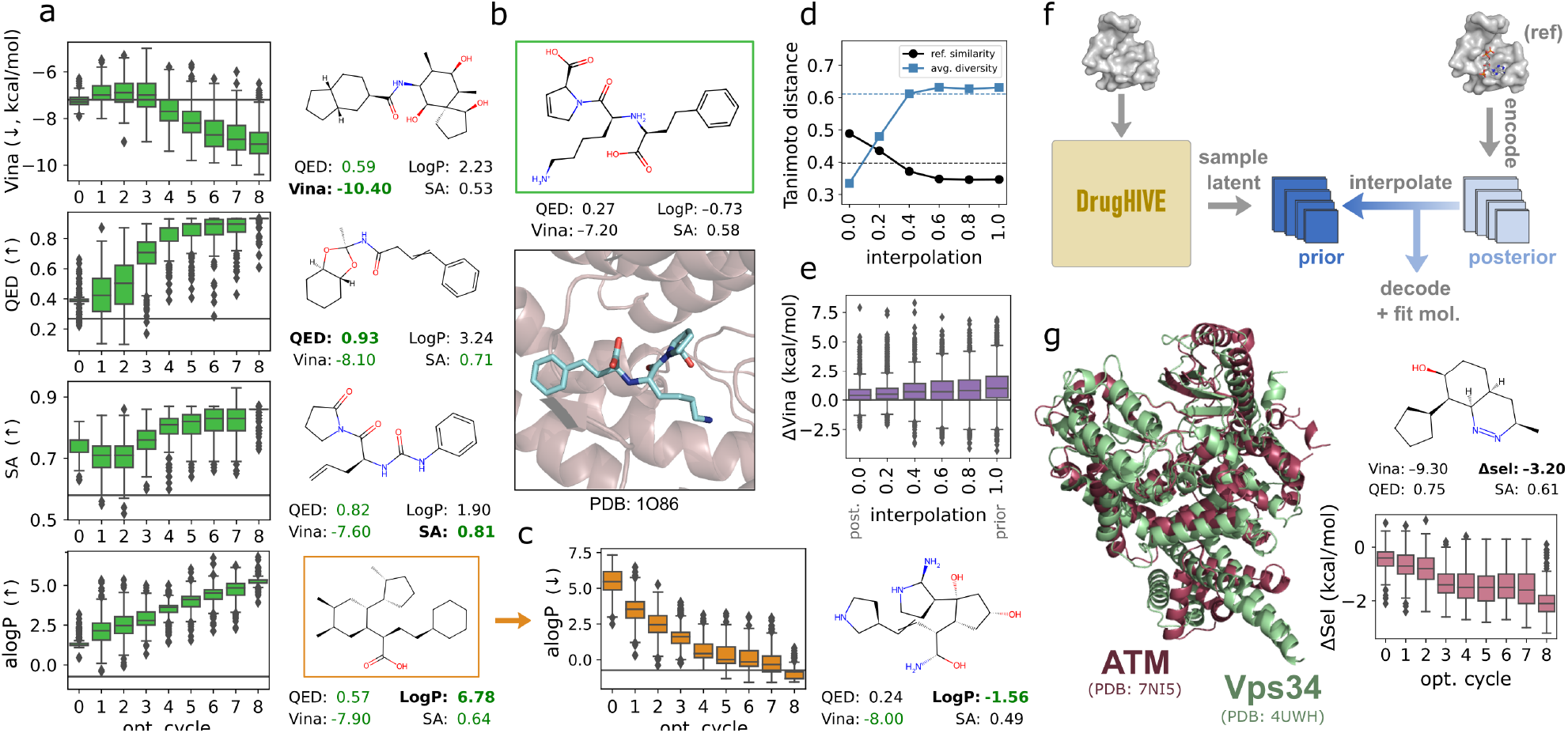
Prior-posterior sampling and multi-objective evolutionary optimization. **(a)** Optimization of the drug molecule Lisinopril bound to the human angiotensin converting enzyme (ACE) receptor (PDB ID 1O86). (left) Boxplots showing the distributions of generated molecules at each optimization cycle. Each boxplot represents a distribution of 400 molecules. (right) Best molecule from the optimization procedure and reported properties, with green text indicating successful improvement. **(b)** Initial molecule (Lisinopril) for the optimization process. **(c)** Optimization of hydrophobicity (alogP) in the downward direction. Starting point was the best molecule from the previous (upward) hydrophobicity optimization. **(d)** Plot of average diversity and similarity to reference molecule for molecules generated from a diverse set of 100 PDB structures (test set) at varying prior-posterior interpolation values. The average diversity of the reference ligands in the test set (*blue dashed line)* and the average similarity of reference ligands in the test set (*black dashed line*) are shown. **(e)** The distributions of *ΔVina* scores of generated ligands as a function of interpolation value. *ΔVina* is calculated as the predicted affinity (*Vina*) score of the reference ligand minus that of the generated ligand. **(f)** Schematic of prior-posterior sampling procedure. **(g)** Generation of molecules with optimized selectivity for the human ataxia-telangiectasia mutated (ATM) kinase and against the human vacuolar protein sorting (Vps34) kinase. (left) Superimposed aligned structures of ATM (PDB ID 7NI5) and Vps34 (PDB ID 4UWH) kinases. (top-right) The best molecule from optimization procedure and reported properties. (bottom-right) Boxplots showing the distribution of selectivity (*ΔSel=Vina*_*ATM*_ *– Vina*_*Vps34*_) of generated molecules at each optimization cycle.

In Figure 2a, we show the successful optimization of binding affinity (Vina), drug-likeness (QED), synthesizability (SA), and hydrophobicity (alogP) values. For QED, SA, and alogP, we perform multi-objective optimization by including binding affinity as a secondary target in the fitness function. We see a significant improvement for each target property, including a 44% improvement in predicted binding affinity for the best molecule despite the fact that Lisinopril, an FDA approved ACE inhibitor in humans used to treat hypertension, heart failure, and myocardial infarction, shows strong inhibition of the target receptor, with a reported experimental K_i_=0.27 nM^26^. Further, we successfully increase the drug-likeness from a low value of 0.27 to a high value of 0.93 and a moderate synthetic accessibility score of 0.58 to a high value of 0.82, in both cases while simultaneously improving binding affinity. In Figure 2c, we first optimize the alogP value in the upward direction, beginning with the slightly hydrophilic Lisinopril (alogP=–0.73) and ending with a very hydrophobic molecule (alogP=6.78). We then take the resulting molecule and optimize the alogP value in the downward direction, reversing the process to result in a hydrophilic molecule (alogP=–1.56).

In Figure 2g, we show results from the generation and optimization of a set of ligands for selectivity toward the human ataxia-telangiectasia mutated (ATM) kinase and against the human vacuolar protein sorting (Vps34) kinase. It has recently been shown that ATM kinase is an important potential target for the treatment of Huntington’s disease and that it is important for ATM inhibitors to exhibit high selectivity over Vps34^27^. We use DrugHIVE to generate an initial set of ligands for the ATM receptor (PDB ID 7NI5), which we then optimize for selectivity against the Vps34 receptor (PDB ID 4UWH). The results show considerable improvement of selectivity, with the best molecule having a significantly better binding affinity score for ATM (–9.3 kcal/mol) compared to Vps34 (–6.1 kcal/mol). These experiments demonstrate that the latent encoding learned by DrugHIVE is a powerful representation of chemical space, enabling effective multi-objective optimization of molecular structures with a high degree of control.

### Lead optimization with spatial prior-posterior interpolation

The hierarchical prior structure of DrugHIVE preserves spatial context and allows molecules to be spatially modified by applying prior-posterior sampling on a spatial subset of the latent representation (as shown in Figure 3b). To demonstrate, we apply this method to common drug design problems such as linker design, substructure modification (scaffold hopping), fragment growing, and molecular pattern replacement (Figure 3a).

**Figure 3:**
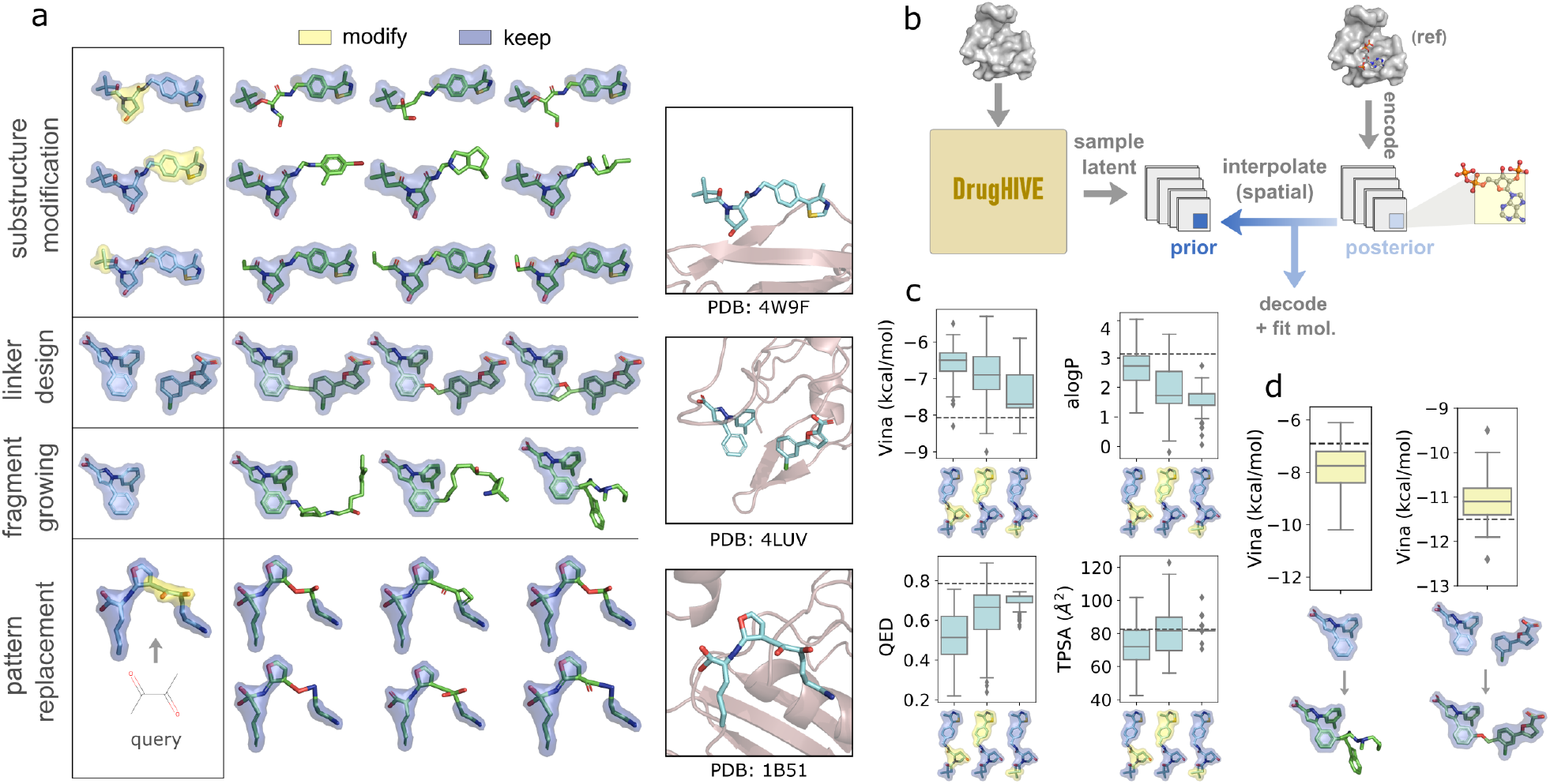
Spatial prior-posterior sampling. **(a)** Examples of substructure optimization (fragment hopping), linker design, fragment growing, and high throughput pattern replacement. Yellow highlighted region indicates the part of the molecule to be modified and blue highlighted region indicates the part to be retained. **(b)** Schematic of spatial prior-posterior sampling procedure. **(c)** Boxplots showing the distributions of properties for each of the modified fragments from the substructure modification experiment with property value of initial molecule shown (*black dashed line*). **(d)** Boxplot showing the distribution of predicted binding affinity (*Vina*) scores for the generated molecules from the fragment growing experiment (left) compared to the *Vina* score of the initial fragment (*black dashed line*) and the linker design experiment (right) compared to the *Vina* score of the linked molecule (PDB ID 1XT) designed by Frank et. al. (*black dashed line*).

#### Linker design

Fragment based screening is a common tool in the early drug discovery toolkit^28^, the result of which is a set of relatively small molecular fragments with modest affinity for the target receptor. Given two or more such fragments, the challenge is to then connect the individual fragments into a coherent, high-affinity drug-like molecule. Ordinarily, this is a manual and time-consuming process, but we show that our model can be used to automatically connect a set of individual fragments as oriented in the cocrystal structure. As an example (Figure 3a), we show that DrugHIVE successfully designs a molecular linker to connect the individual fragments (PDB IDs 1DZ and 1XS) from a screening experiment for replication protein A (PDB ID 4LUV)^29^. All of the resulting ligands have a *Vina* score better than either of the individual fragments, with 21% of generated molecules having equal or better *Vina* score and 51% having equal or better *synthetic accessibility* score compared to the linked molecule (PDB ID 1XU) designed by ref.^29^ which has an experimentally measured k_d_=20 μM.

#### Fragment growing

Another common drug design task is to take a known fragment with modest binding affinity due to its size and grow the structure to improve its affinity for the target. Our model can be used to automatically design new substructures to add to an existing molecular structure. To demonstrate, we show that DrugHIVE successfully grows one of the fragments (PDB ID 1DZ) from the same fragment screening experiment, significantly increasing its predicted binding affinity for the target receptor (Figure 3d), with 85% of generated molecules having an improved *Vina* score.

#### Substructure modification (scaffold hopping)

A common drug optimization technique is the systematic replacement of either chemical scaffolds or R-groups on a candidate molecule in order to improve its properties or grow a candidate set^30^. Starting with the ligand (PDB ID 3JU) bound to the von-Hippel Lindau (VHL) tumor suppressor (PDB ID 4W9F), we show how automatic substructure optimization can be achieved using spatial prior-posterior sampling. We apply BRICS^31^ decomposition to break the ligand, an agonist with reported K_d_ =3.27 µM^32^, into molecular fragments. For each fragment, we generate a set of molecules for which we only modify the fragment substructure, leaving the rest of the molecule unchanged (Figure 3a). For each of the BRICS fragments, we then analyze the distribution of properties for the generated molecules to learn the contribution of each fragment to each particular property. In Figure 3c we show that, depending on which fragment is modified, there are significant differences in the effect on each of the properties (*ΔVina*, alogP, TPSA, QED). For instance, we observe that drug-likeness scores are improved only for fragment 2, and that modification of fragment 3 leads to the strongest decrease in hydrophobicity (alogP) values while having almost no effect on topological polar surface area (TPSA). We also see that modification of fragment 1 leads to the strongest increase in *Vina* score, while fragment 3 has relatively little effect. This approach can generate valuable insights for lead optimization and structure-activity relationship (SAR) analysis.

#### Pattern replacement

Given a molecular SMILES or SMARTS pattern, we can modify the molecular structure(s) corresponding to that pattern for a single molecule or a set of molecules, using DrugHIVE to generate a new set of molecules with the input pattern replaced. This functionality could be used to modify fragments with known toxic, promiscuous, or otherwise undesirable properties in existing or newly generated drug collections, mitigating the need to screen out candidates with otherwise promising qualities. One obvious application of this is in the replacement of PAINS patterns, which are a set of molecular substructures shown to be correlated with false positive results in HTS assays^33^. It should be noted that these molecular patterns do not always lead to experimental interference and therefore screening decisions must be taken with some care^34^. To avoid both the risks of false positive experimental results and false negative screening decisions, we can simply replace all PAINS patterns in an existing collection using generative spatial modification. As an example, we apply this capability to modify all PAINS structures in a set of molecules previously generated for the peptide binding protein OppA (PDB ID 1B51). In Figure 3a, we show a collection of molecules resulting from replacing one query pattern. Further, this pattern replacement capability is scalable to large collections of molecules, as the run time per molecule is on par with fast virtual docking algorithms.

### Drug design for predicted target structures

A major limitation of SBDD methods is the reliance on a high-resolution crystal structure of the target receptor. Protein crystal data is growing rapidly, with the number of deposited structures to the PDB averaging over 9,000 per year in the last decade^35^. Still, less than 20% of the human proteome is currently resolved^36^. Recently, the success of sequence-to-structure prediction models such as AlphaFold2^12^ have provided high accuracy structures for previously unknown proteins, already filling in a majority of the missing structural data for foldable regions of the human proteome with high confidence structures^36^. We show that DrugHIVE is able to overcome this limitation by generating high affinity molecules for receptors using AlphaFold predicted structures. To test the effectiveness of using predicted structures for generative drug design, we generate ligands for a test set of target receptors for which we have both the AF prediction and the PDB crystal structure. The process for AF target curation is outlined in Figure 4a. It is important to stress that no crystal structure alignment information was used in selecting the AF targets. Filtering is based solely on sequence alignment, to ensure adequate sequence similarity, and AF confidence score (pLDDT), to ensure high confidence predictions.

**Figure 4:**
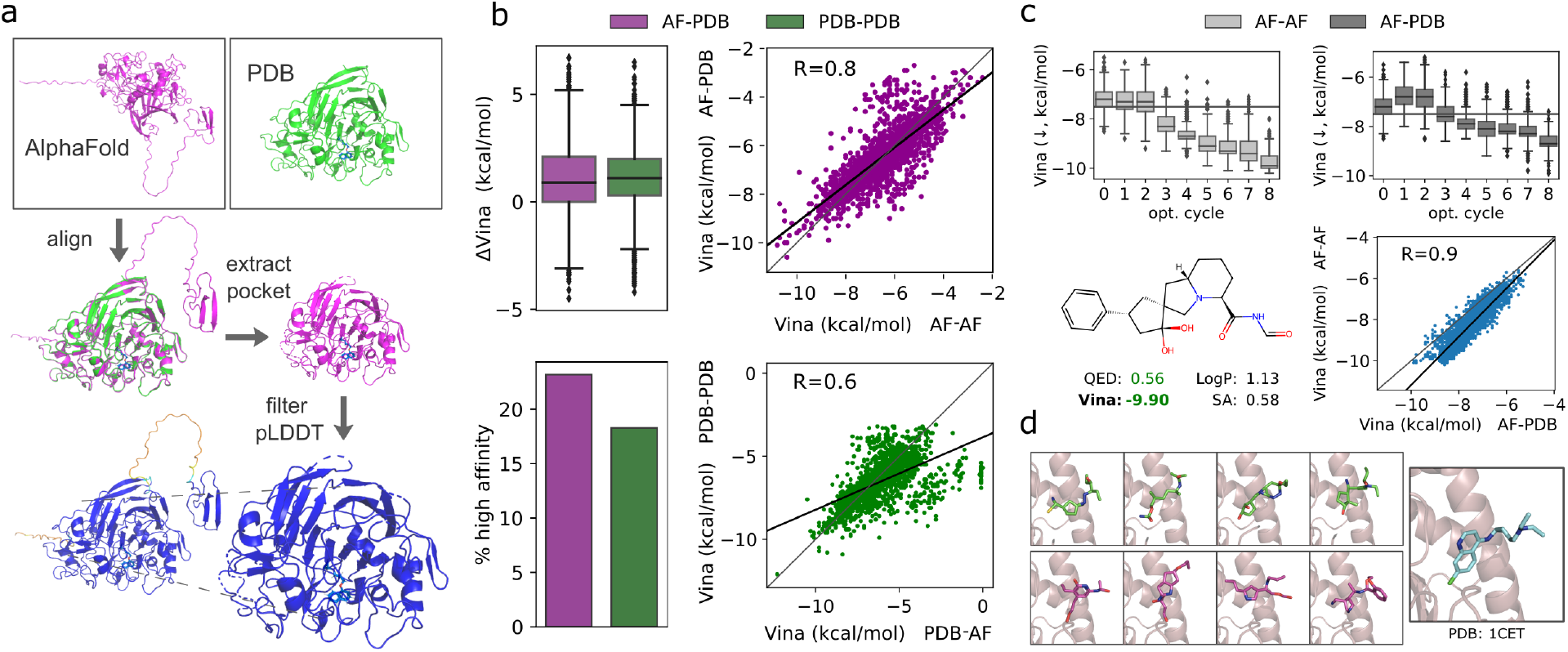
Molecular generation using AlphaFold predicted structures. **(a)** Overview of the pipeline for creating the dataset of AlphaFold (AF) test structures. **(b)** Comparison of AF-generated molecules with crystal structure (PDB) generated molecules. (left) AF-generated molecules receive slightly better predicted binding affinity scores compared to PDB-generated molecules, with a larger portion of high affinity molecules (predicted affinity better than crystal ligand). (right) Scatterplots showing significant correlation of predicted *Vina* scores between molecules docked to AF and PDB structures. The AF-generated molecules have a slightly higher Pearson correlation (*R*=0.8) compared to the PDB-generated molecules (*R*=0.6). **(c)** Evolutionary molecular optimization of the drug Lisinopril bound to human angiotensin converting enzyme (ACE) receptor (PDB ID 1O86) using the AF-predicted receptor structure. (top) Boxplots showing the distributions of *Vina* score for molecules at each step of the optimization process docked to the AF and PDB receptors. (bottom-left) The best molecule from optimization process along with reported properties. The predicted binding affinity for the PDB receptor is shown. (bottom-right) Scatterplot showing a high correlation (Pearson *R*=0.9) between AF and PDB docking scores for all molecules from the optimization process. **(d)** Random examples of molecules generated for PDB ID 1CET using the PDB receptor (green) and the AF receptor (magenta). All molecules are displayed in the PDB receptor.

We dock the AF-generated ligands into both the AF and crystal receptors and compare the binding affinity predictions to ligands generated for the PDB receptor. In Figure 4b, we show the distribution *ΔVina* scores for the AF and PDB generated ligands docked to corresponding PDB receptors. The range of affinities is very similar between the two sets, with a slightly better median value of the AF-generated ligands. The proportion of ligands with a better docking score than the crystal ligand is around 20% for both sets, and slightly better for the AF generated ligands. We also show that there is good correlation between the Vina scores when docked to both the PDB and AF receptors for AF-generated ligands (*R*=0.8) and PDB-generated ligands (*R*=0.6).

Further, we show in Figure 4c that DrugHIVE can optimize the predicted affinity of ligands using the AF pocket. We run the optimization process beginning with Lisinopril placed in the AF predicted structure corresponding to PDB ID 1O86, resulting in a molecule with a similarly high predicted binding affinity compared to the same optimization process using the PDB pocket (Figure 2a). We then docked all molecules generated during the optimization process into the PDB pocket and calculated the Pearson correlation coefficient for the two sets of affinity scores. The correlation is high (*R*=0.8), with the AF-docked scores consistently underestimating the PDB-docked scores— a bias that is exaggerated for higher affinity ligands. Despite this underestimation, the high correlation allows for the DrugHIVE optimization process to significantly improve binding affinity scores of the initial ligand.

## DISCUSSION

We introduce DrugHIVE, a new approach to the structure-based design of drug-like molecules based on a deep hierarchical variational autoencoder architecture. Our motivation in designing our model was based on the observation that, even though molecular systems clearly possess structure and properties at varying spatial scales, previous encoder-decoder models have failed to adequately capture these relationships. Our model’s hierarchical prior enables the model to better learn the distribution of intra- and inter-molecular relationships and generate high affinity drug-like molecules with an unprecedented degree of spatial control. In particular, DrugHIVE displays an impressive ability to optimize the properties of molecules over a wide range of values within the context of a protein receptor by performing local exploration of the latent space.

Beyond finding a more natural representation of chemical space, our second motivation in designing the model was improving practical applicability for drug-design tasks. Many tasks in early drug design are time consuming and rely significantly on the intuition and expertise of researchers. While such expert knowledge will always be a crucial part of the process, generative models can help speed up cycle times by automating the de-novo design and modification of molecular structures. In this work, we show how DrugHIVE can be used to perform rapid and effective linker design, scaffold hopping, fragment growing, and high throughput pattern replacement—all of which ordinarily require careful expert attention. Currently, DrugHIVE can only generate molecules that fit within the input density grid, which limits its applicability to larger molecules. The grid width that we chose (24 Å) fits most drug-like molecules, but extending this approach to larger structures quickly grows the computational requirements. Our dual-resolution implementation mitigates this issue significantly by lowering the memory requirements, but grids much larger than 48 Å would still require expensive multi-GPU training. Although we think that the space-filling property of the grid-based representation has significant advantages, other generative methods such as graph and point-cloud based models are being developed to avoid these limitations^20–22,37^.

DrugHIVE demonstrates that generative models provide an exciting new avenue for expanding our reach into the vast drug-like chemical space, the majority of which is unexplored and inaccessible. For the 100 diverse protein structures in our test dataset, 10.3% of the molecules generated by DrugHIVE have a predicted affinity higher than that of the crystal ligand—significantly higher value than the 4.6% observed for a randomly chosen ZINC screening set. This means that performing hit identification by screening generated molecules already surpasses the efficiency of current screening datasets and will surely improve as the field progresses. Additionally, virtually none of the generated molecules exist in the training dataset, which means that DrugHIVE is able to access previously unexplored regions of chemical space.

The synthesizability of generated molecules does appear to be lower than that of current screening libraries, which is a significant factor in successful hit identification. This is partly due to the relatively low synthesizability of crystal ligands compared to that of drug-like molecule collections— especially those collections created from chemical synthesis enumeration such as Enamine REAL^4^. However, there are promising current research efforts to help overcome this limitation. Future work to improve synthesizability of generated molecules, such as synthon based generation^16,38^, could significantly improve their practical utility for hit identification. Improvements to retrosynthesis planning using deep learning methods are leading to more accurate identification of synthesizable compounds^39,40^, and advances in chemical synthesis methods are expanding the accessible molecular space^41–43^.

DrugHIVE is not limited to existing crystal structures. Even though recent work has called into question the utility of using AF predicted structures in drug design applications^44^, there have also been examples of where its use has been successful^45^. our experiments show that molecular generation with DrugHIVE using AF structures does generalize well to the PDB structures we tested. We show that over 20% of ligands generated for a diverse set of AF-predicted structures have higher predicted affinities than the crystal ligand when docked to the PDB crystal structure, with a high correlation between docking scores (*R*=0.8). Further, we show that optimization of binding affinity using the AF predicted structure yields molecules with significantly improved predicted binding affinities for the PDB structure. This means in the case where the crystal structure is not available, docking to a high-confidence AF structure should give a reasonable estimate of the crystal structure docking score—though not necessarily the correct binding pose. The implications are significant given that 58% of the human proteome residues, and perhaps all of the remaining foldable regions without crystal data, are currently predicted with high confidence by AlphaFold2^36,46^. Finally, it is important to note that the application of our model is agnostic to the choice of virtual docking algorithm, and as virtual docking methods improve so will the performance of DrugHIVE in many drug design tasks. Such progress may be forthcoming with efforts to design deep learning models that learn protein-biomolecule co-folding beginning to show promising results^47–49^.

By developing a model with more fine-grained control over molecular generation, we hope to accelerate the impact of generative models on early drug discovery. We help demonstrate the great potential that generative models have for automating a wide array of drug design tasks. Further progress and widespread adoption of these methods has the potential to significantly accelerate and lower the cost of pre-clinical drug development. With an estimated cost of $0.04 to $0.20 per thousand generated molecules at current cloud computing rates, on par with current high-throughput virtual screening methods, DrugHIVE is a powerful and scalable tool for automated structure-based drug design.

## METHODS

### Model

The DrugHIVE model is illustrated in Figure 1. Input molecules are first converted to molecular density grids, where each atom is represented by a multi-channel pseudo-Gaussian density and each channel represents a different atomic feature. For each atom, an identical atomic density is placed in each grid channel corresponding to its features—similar to ref.^19^. In this work, we use a subset of 15 feature channels: nine atomic elements (C, N, O, F, P, S, Cl, Br, I) and six other atomic features (H-bond donor, H-bond acceptor, aromaticity, positive charge, neutral charge, negative charge). For evaluation, we use a version of our model with the nine atomic elements only. The molecular grid is input into the network via two pathways (Figure 1c). A single-channel full resolution grid, representing atomic occupancy, is input into the top of the network. All other atomic feature channels are downsampled by a factor of two and input into the network after the first downsampling layer. Model outputs mirror this arrangement. By segregating occupancy from atomic features in this fashion, we retain atom position information while significantly decreasing grid sparsity and memory load.

The DrugHIVE model (Figure 1d) is based on the hierarchical variational autoencoder (HVAE) architecture^50–52^. The goal is to learn the distribution of ligand density conditioned on receptor density from the dataset. However, unlike for a standard VAE, the latent representation of an HAVE naturally retains spatial context (Figure 1b). Therefore, we have a generative model of the form *p*(*D*_*lig*_, *D*_*rec*_, ***z***) = *p*(***z***)*p*(*D*_*lig*_ |***z***, *D*_*rec*_), where *p*(*D*_*lig*_ |***z***, *D*_*rec*_) is the likelihood function (decoder), the prior is represented as *p*(***z***) = ∏_*i*_ *p*(*z*_*i*_|*z*_<*i*_) and the approximate posterior as *q*(***z***|*D*_*lig*_, *D*_*rec*_) = ∏_*i*_ *q*(*z*_*i*_ |*z*_<*i*_, *D*_*lig*_, *D*_*rec*_). We represent the likelihood function and approximate posterior as neural networks *p*_*θ*_ and *q*_*ϕ*_, respectively. The latent representation is a set of variables partitioned into a set of disjoint groups ***z*** = {*z*_1_, …, *z*_*n*_} where each group is conditioned on the lower groups in the hierarchy, *z*_*i*_ ∼ *q*(*z*_*i*_ |*D*_*lig*_, *D*_*rec*_, *z*_<*i*_). Because the individual latent variables retain spatial context at varying resolutions (Figure 1b), we hypothesize that the structure of the latent space is a better reflection of molecular structure space.

### Pseudo-Gaussian density

We represent atoms as pseudo-Gaussian densities by first encoding them to a grid using what we call center-of-mass (COM) encoding followed by a Gaussian convolution. We consider each atomic coordinate *c* ∈ ℝ^3^ to represent the center of a cube with the same dimensions as a single voxel. Then, to encode an atom to the grid, we calculate the proportion of the cube contained in each grid voxel and add that value to the corresponding voxel value. Therefore, to acquire the COM encoding of a set of *N* atoms with grid coordinates *c*_*j*_ ∈ {*c*_1_, …, *c*_*N*_} on a grid with spacing *a* = 1, we add up each atom’s contribution to each grid voxel *g*_*i*_ ∈ *G*_*COM*_,

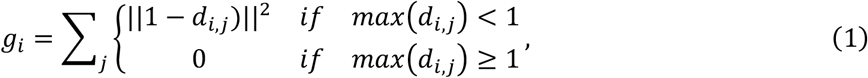

where c_gi_ are the grid coordinates of the *i*^*th*^ voxel, *c*_*j*_ are the grid coordinates of the *j*^*th*^ atom, and *d*_*i,j*_ = |*c*_*gi*_ − *c*_*j*_|. Importantly, precise positional information is retained and remains recoverable after the encoding process. Given a set of adjacent grid points containing an encoded atom 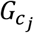, the original coordinate *c*_*i*_ can be exactly recovered using the center of mass equation,

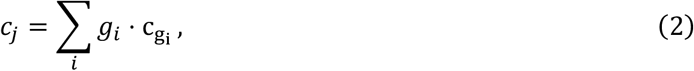

where the value of *g*_*i*_ represents the mass of each grid point. This is in contrast to the simple one-hot encoding scheme where each atom is represented by a single voxel and positional precision is lost.

Next, we perform a convolution of the COM grid with 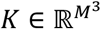, a Gaussian kernel with side length *M*, to get the density grid *D* = *K* * *G*_*COM*_. The value at each kernel point *k*_*i*_ ∈ *K* is calculated according to,

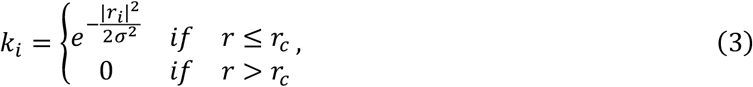

where *r*_*i*_ is the distance of the grid point from the grid center and *r*_*c*_ is a cutoff radius. The resulting density grid is an approximation of an exact Gaussian encoding of each atom and can be summarized as the Gaussian blurring or diffusion of the initial COM encoding pattern. The advantage of this procedure is that the initial COM encoding is very fast to compute, and the Gaussian convolution operation can be represented as a standard convolutional neural network layer with a fixed Gaussian kernel. The choice of convolutional kernel is customizable, and the whole process can be easily and efficiently implemented using common Python scientific and machine learning libraries.

### Atom fitting

The output of our model is a molecular density grid which must then be mapped to a molecular structure (Figure 1e). To achieve this, we employ custom atom-fitting and bond-fitting algorithms. The atom-fitting algorithm first iteratively places atomic coordinates within the grid with the goal of minimizing the residual density,

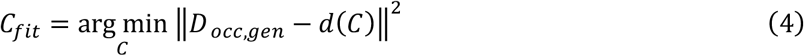

where *D*_*occ,gen*_ is the predicted atomic occupancy density and *d* is the mapping from coordinates to density grid *D*. Next, atomic features are assigned to each atom using the density in the upsampled atomic feature grid. Atomic number is assigned to the highest density value across all atomic number channels at each atom location. The remaining atomic features are assigned based on a density threshold value, with disputes between conflicting features (e.g., formal charge ± 1) resolved by selecting the feature with the highest density value. Next, bonds are added based on the geometry and features of the atom set, similar to ref.^19^, to create a molecular structure.

### Training

The model is trained to minimize the objective,

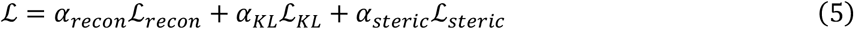

where ℒ_*recon*_ is the reconstruction loss, ℒ_*KL*_ is the Kullback-Leibler (KL) divergence between the prior and posterior distributions, ℒ_*steric*_ is the loss for steric clash between the ligand and receptor density, and each *α* is a scaling term.

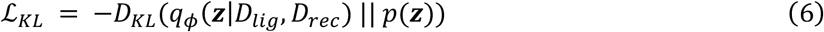

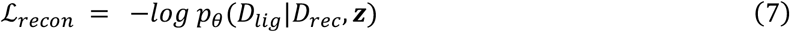

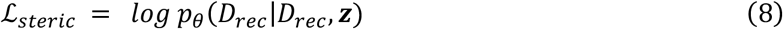

Because the grid representation is sparse, we apply a mask to scale the reconstruction loss for zero grid values in order to focus the loss signal on non-empty grid locations. We achieve this by scaling the loss for zero grid values by a factor,

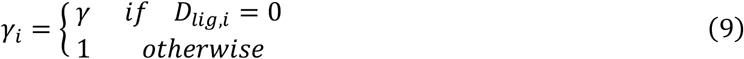

yielding the scaled reconstruction loss,

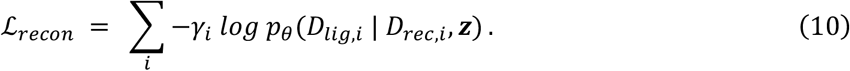

### Data

We use two datasets in the training and evaluation of DrugHIVE, PDBbind (v2020, refined set)^25^ and a subsample of the ZINC drug-like subset^5^. We chose the PDBbind refined set as it provides a set of high-quality crystal structures of ligands bound to protein receptors. We remove structures with ligands that are too wide to fit well into the density grid (>16 Å) or that have more than a threshold number of atoms (>43). A test set of 100 receptor structures was randomly selected from the dataset. We also train on subsample of the ZINC dataset as unbound drug molecules to improve training and reduce overfitting to the relatively small number of example ligands in the PDBbind dataset. To acquire the subsample, we start with the ZINC drug-like subset and sample about 27 million molecules uniformly by alogP and number of heavy atoms, filtering out the small number of structures with heavy atoms not in our atomic element set (Z ∈ {C, N, O, F, P, S, Cl, Br, I}).

For evaluation of generating from AlphaFold predicted structures, we use a dataset of 100 AF structures for which we also have the PDB structure in our dataset. To create this dataset, we first identify all structures from the PDBbind dataset where the receptor pocket is comprised of a single protein chain and that chain is available from the AlphaFold Protein Structure Database^53^. We then perform sequence and structural alignment of the AF structure to the PDB structure using PyMOL *align* function^54^, filtering out receptor structures with less than 70% sequence identity. We then extract the binding pocket by cutting off any excess residues based on sequence alignment and remove all residues with atoms > 24 Å from the ligand. We then filter out any structures without a high confidence score (pLDDT < 70) for any residue within 5 Å of the ligand. Finally, we randomly sample 100 of the remaining receptors for the AF test set.

### Sampling

DrugHIVE has multiple sampling modes. Once a model is trained, new molecular structures can be generated by inputting a protein structure into the protein encoder and a randomly sampled set of values from the prior into the ligand decoder. We call this mode *prior sampling*, and the result is a randomly generated density that can then be fit with a molecular structure. Since known ligands can be encoded into a set of latent values, we can repeat the same procedure using the latent values from a previously encoded ligand structure instead of randomly sampling from the prior. We call this *posterior sampling*, and the result is an output density very similar to the true input ligand density.

We combine these two sampling modes into what we call *prior-posterior sampling (*Figure 2f) to generate densities of varying similarity to an initial ligand density. To achieve this, we repeat the procedure for posterior sampling while also sampling a random set of values from the latent prior. Starting with the values sampled from the posterior, we then interpolate toward the values sampled from the prior:

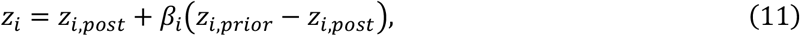

with *β*_*i*_ being the interpolation factor. The latent representation has multiple scales, and one can specify a different interpolation factor for each scale. Because different molecular properties are encoded at different scales (Figure S4), this leads to a high degree of control over the variation of molecular details in generating new structures.

The latent space of DrugHIVE is comprised of a set of tensors, each representing information at a specific resolution of molecular space (Figure 1b). This varying resolution is the result of downsampling layers in the neural network. In our model each latent tensor retains spatial context, so we can do *spatial prior-posterior sampling* by performing prior-posterior sampling over a subset of the density grid (Figure 3b). For this sampling mode, we choose a region of the input grid on which to perform prior-posterior sampling. For each latent tensor, we determine which values correspond to the modification region and which fall outside by mapping the tensor values back to the full resolution grid. We then perform prior-posterior sampling, setting the interpolation factors equal to zero for all latent values outside of the modification region. The result is a generated structure similar to the input ligand except for within the modification region, which will differ more or less according to interpolation factor value.

Another means of controlling the generation process is through a scaling (temperature) factor on the variance of each latent variable. The DrugHIVE latent prior is a high dimensional normal distribution, where each latent value is sampled from a distribution, *z*_*i*_∼*N*(*µ*_*i*_, *σ*_*i*_*τ*_*i*_), where *µ*_*i*_ is the mean, *σ*_*i*_ is the variance and *τ*_*i*_ is the temperature factor. As with the prior-posterior interpolation factor, the temperature factor can be varied across latent scales and spatial subsets. The default temperature factor during training is *τ*_*i*_ = 1. Generally speaking, decreasing the temperature factor has the effect of lowering the variability of generated samples^51,52^.

### Evolutionary molecular optimization

Evolutionary (genetic) algorithms are employed in a wide range of multi-objective non-gradient optimization problems and have a long history of use in drug discovery^55^. They mimic the process of biological evolution to effectively search through spaces of high complexity where ordinary search methods would be prohibitively inefficient or difficult to implement. We apply an evolutionary algorithm coupled with prior-posterior sampling to search the latent space of DrugHIVE and optimize molecular properties. In our case, the genetic information is represented by the latent encoding of molecules and the mutation rate is represented by the prior-posterior interpolation factor (*β*).

#### Initialization step

Generate an initial population of *N*_*c*_ molecules. Apply the fitness function and cluster the molecules (Tanimoto similarity < 0.2), keeping the most fit individual from each cluster. Select the *N*_*p*_ best molecules from the current population as “parents”.

#### Optimization step (repeat until converged)

Generate a new population of *N*_*c*_ “child” molecules from parents. Apply the fitness function and cluster, keeping the most fit individual from each cluster. Replace least fit parents with child molecules having better fitness score.

The fitness function will vary based on desired outcomes but, in general, will be some measure of distance between the observed molecular property value(s) and the desired ones. For example, in the case of binding affinity optimization, the predicted binding affinity score is used as the fitness measure. For multi-objective optimization, the fitness function can be a combination of (e.g., weighted sum) of individual fitness scores. In our non-affinity property optimization experiments, we seek to maintain predicted binding affinity while optimizing the targeted property by removing all child molecules from the population with a binding affinity score worse than a threshold value.

### Force field optimization

Prior to docking, we perform force field optimization on all molecular structures using the MMFF94 force field^56^ in RDKit. Unlike some other groups, we do not use the Universal Force Field^57^ due to a lack of robustness in convergence that we empirically observed in this work. We also do not relax structures in the context of the receptor for a couple of important reasons. The first reason for this is that docking algorithms are designed to take as input an energy minimized ligand structure. For docking algorithms based on AutoDock, it is important to input chemically valid molecular coordinates^58,59^. In fact, even ligands from crystal structures that are geometrically distorted by the receptor are recommended to have their structures optimized before proceeding to docking^58^. Second, convergence to an optimal minimum becomes less reliable as the size of the system grows. Including the surrounding receptor atoms in the calculation, even if fixed to their initial positions, may lead to a higher likelihood of convergence failure, false positives, or geometric distortions outside of the distribution of structures on which the parameters of the Vina scoring function were fit^60^. This step is especially important with de-novo generated structures, which are likely to be far from an energy minimum. Using unconverged structures or, even worse, choosing to skip force field optimization altogether on structures before docking could lead to significantly exaggerated docking scores (Figure S3).

### Evaluation properties and metrics

Binding affinity is estimated for each ligand with the *Vina* docking score calculated with QuickVina 2^61^, a speed optimized version of AutoDock Vina. We define *ΔVina* as the difference between the Vina score of a generated ligand and the reference ligand (e.g., the ligand from the crystal structure). Molecular *similarity* is calculated as one minus the Tanimoto distance between the molecular fingerprints of a pair of molecules. Molecular *diversity* is defined on a set of molecules as the average Tanimoto distance between each pair in the set. The Quantitative Estimate of Drug-Likeness (*QED*) score estimates the drug-likeness of a molecule by combining a set of molecular properties^62^. The Synthetic Accessibility (*SA)* score estimates the ease of synthesis of a molecule^63^.

### Evaluation

We evaluate the molecules generated by our model against competitors and a set of randomly sampled molecules from the ZINC20 drug-like subset by comparing the predicted binding affinity of the molecules. We generate 200 molecules for each of the receptors in the test set for each of the methods and calculate QED, SA, and average diversity using RDKit. We then perform force field optimization on each of the molecules using the MMFF94 force field in RDKit. We virtually dock each molecule to the corresponding receptor using QuickVina 2 (with default settings and a 20 Å wide bounding box) to predict the binding affinity. However, the Vina score is strongly correlated with number of atoms (Figure S2) in the molecule which prevents direct comparison between sets of molecules with different size distributions. To overcome this, we use bootstrap sampling to estimate the average *Vina* score given a uniform distribution of molecular sizes. First, we bin the generated molecules by size, *n*_*a*_ = *number of heavy atoms*. We choose the largest common interval of molecular sizes over which to sample such that all sets have at least 100 molecules in each bin. This resulted in the interval *n*_*a*_ ∈ [13, 25]. We then randomly sample 20 molecules from each group, aggregate, and compute the average binding score,ρ_*i*_. We repeat this process *N* = 200 times and compute the cumulative average 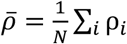. We report this cumulative average for each method as *Vina-n* in Table S1. In Figure S2, we plot the distributions of molecular size and the predicted affinity vs. number of heavy atoms for each generated set.

## ACKNOWLEDGEMENTS

The authors acknowledge helpful discussions with other members of the Rohs lab, including Jared M. Sagendorf. This work was supported by the National Institutes of Health [grant R35GM130376 to R.R.] and Human Frontier Science Program [grant RGP0021/2018 to R.R.].

