## Supplementary Figures S1-S9 and Table S1 for "DrugHIVE: Target-specific spatial drug design and optimization with a hierarchical generative model"

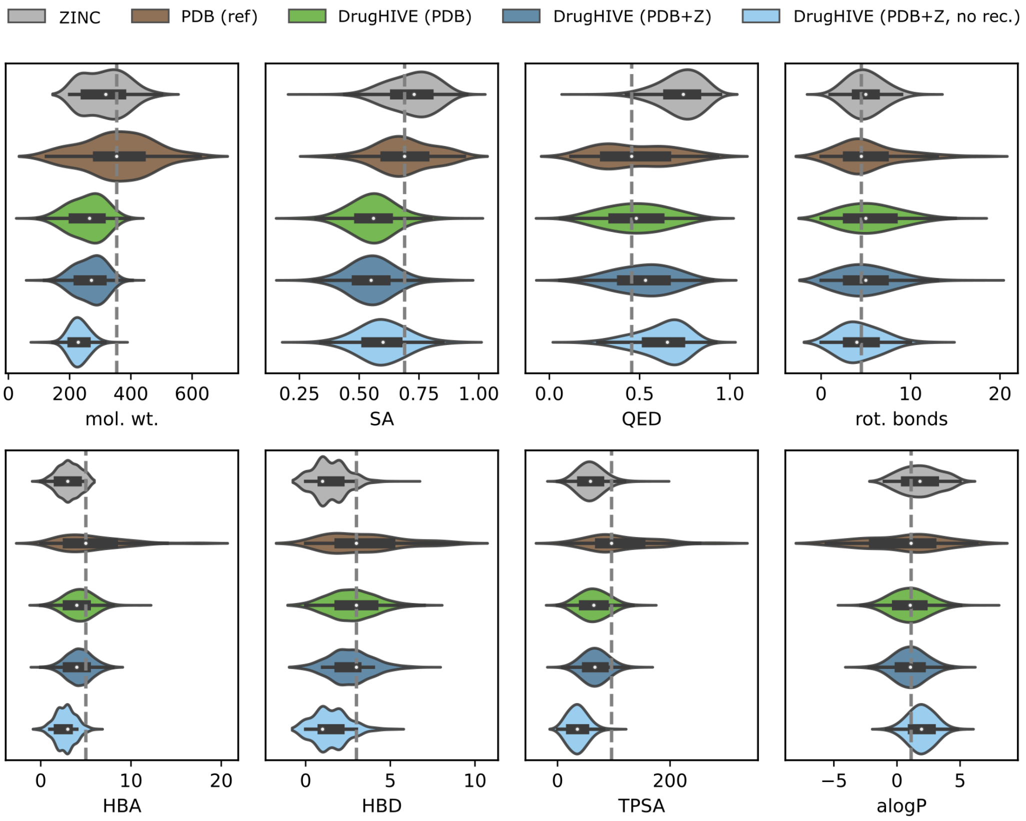


**Figure S1: Property distributions of generated molecules.** Property distributions of generated molecules for different versions of the DrugHIVE model, trained using either PDBbind dataset (PDB), ZINC dataset (Z), or both (PDB+Z), compared to a random subsample of the ZINC dataset and the crystal ligands from the (PDBbind) test set. Molecules generated without a receptor are labeled with no rec. (*mol. wt.: molecular weight, SA: synthetic accessibility, QED: drug-likeness, rot. bonds: rotatable bonds, HBD: H-bond donors, HBA: H-bond acceptors, TPSA: topological polar surface area, alogP: hydrophobicity*)

**Table S1: Evaluation of competing models.** Evaluation of molecules generated for a diverse test set of 100 target receptors chosen from the PDBbind dataset. Summary values (mean ± std. dev./std. err.) are reported for competing models, a random sample from the ZINC drug-like subset, and the test set crystal ligands (Ref). *Vina-n* is the normalized *Vina* score. Diversity is reported as the average per-target molecular diversity. No diversity score is reported for the reference (Ref) set as there is only one molecule per target.

| | Vina-n (↓) | QED (↑) | SA (↑) | Diversity |
|---|---|---|---|---|
| Ref | -7.97±1.98* | 0.46±0.19 | 0.60±0.21 | — |
| ZINC | -6.46±0.06 | 0.72±0.12 | 0.71±0.11 | 0.66±0.02 |
| LiGAN | -6.73±0.06 | 0.46±0.16 | 0.58±0.12 | 0.67±0.02 |
| DiffSBDD | -6.83±0.07 | 0.56±0.17 | 0.68±0.13 | 0.69±0.03 |
| Pocket2Mol | -7.04±0.07 | 0.58±0.19 | **0.71±0.13** | 0.63±0.04 |
| DrugHIVE | **-7.08±0.07** | **0.59±0.15** | 0.56±0.11 | 0.62±0.02 |

*For crystal ligands (Ref), *Vina* mean and std. dev. are reported instead of *Vina-n* mean and std. err.

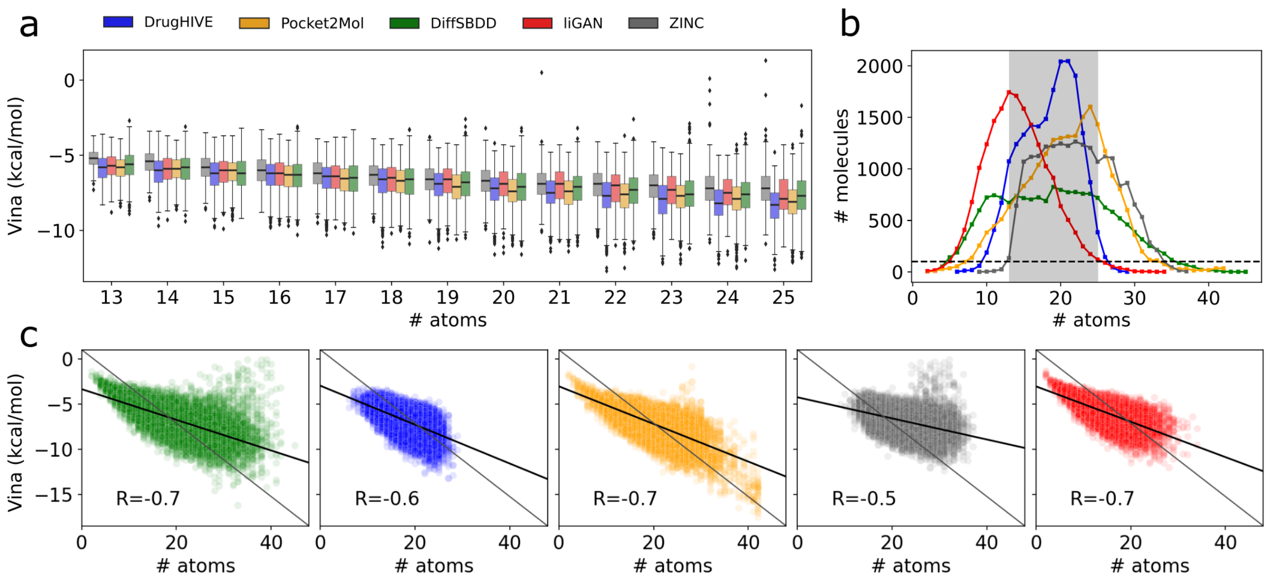


**Figure S2: Binding affinity vs. number of atoms.** Analysis of binding affinity (Vina) score plotted as a function of number of atoms for each of the competing models in Table S1. A random subset of the ZINC dataset is also included for reference. **(a)** Boxplots showing the distributions of Vina score vs. number of atoms. **(b)** Distribution of generated molecule size for each model with cutoff (black dashed line) and test interval (shaded). **(c)** Scatterplots showing the relationship between Vina score and molecule size for each model, with Pearson correlation reported.

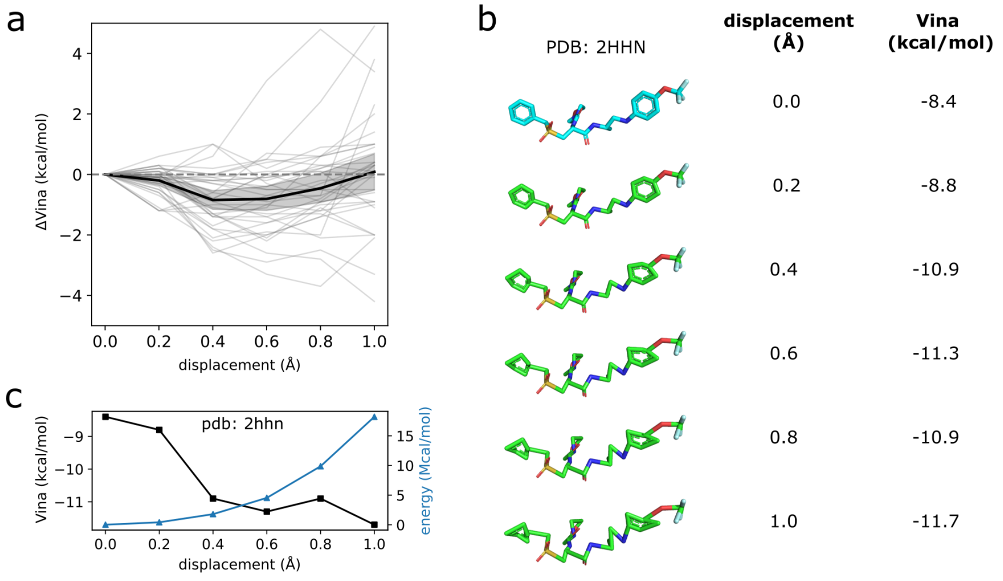


**Figure S3: Force field optimization and predicted affinity score.** Starting with a randomly chosen set of ligands (*n*=33) from the test dataset, we displace each carbon atom by 1 Å in a random direction in increments of 0.2 Å. We then virtually dock each resulting molecule into its receptor using QuickVina 2. **(a)** Relative change in binding affinity score (*ΔVina*) as a function of random displacement. The mean (black line) and 95% confidence interval (grey shadow) are also shown. On average, predicted binding affinity score improves with distortion up to about 0.5 Å (*ΔVina* ≈ -0.84 kcal/mol). **(b)** Distortion of crystal ligand from PDB ID 2HHN along with predicted affinity (Vina) score values. **(c)** Plot of predicted affinity (Vina) score and internal energy (computed with MMFF94 force field) as a function of random displacement for example crystal ligand (PDB ID 2HHN).

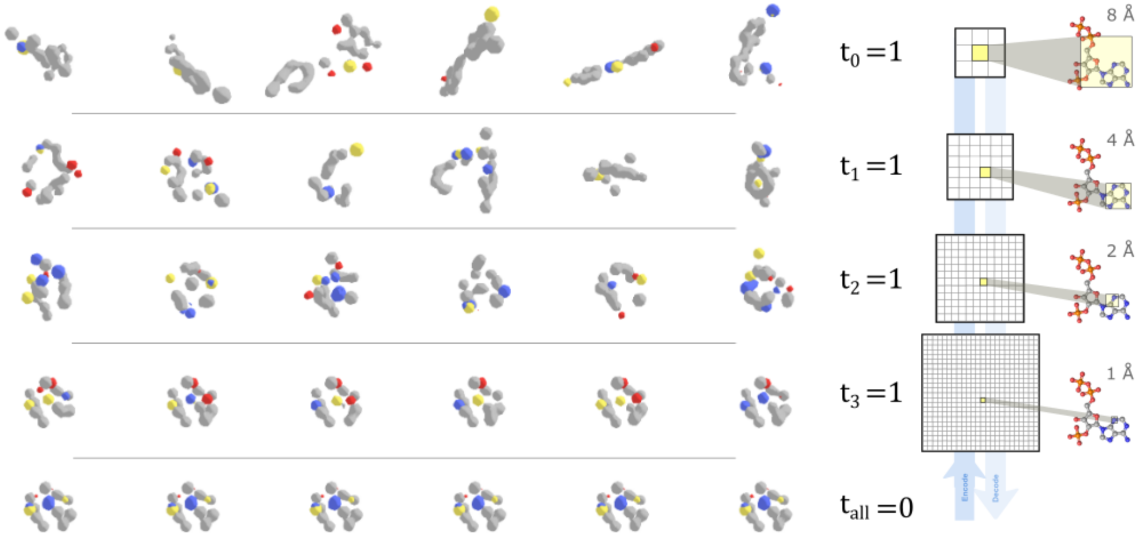


**Figure S4: Latent representation and molecular properties.** Molecular densities generated from DrugHIVE with varying temperature factor settings. Densities are shown as iso-surfaces colored by atomic number channel. (left) Randomly generated densities with all temperature factors set to zero except for the $j^{th}$ latent resolution indicated on the right ($t_j$=1). Bottom row shows case where all temperature factors are set to zero ($t_i$=0, $\forall i$). (right) A schematic depicting the latent resolution of nonzero temperature factor for each row.

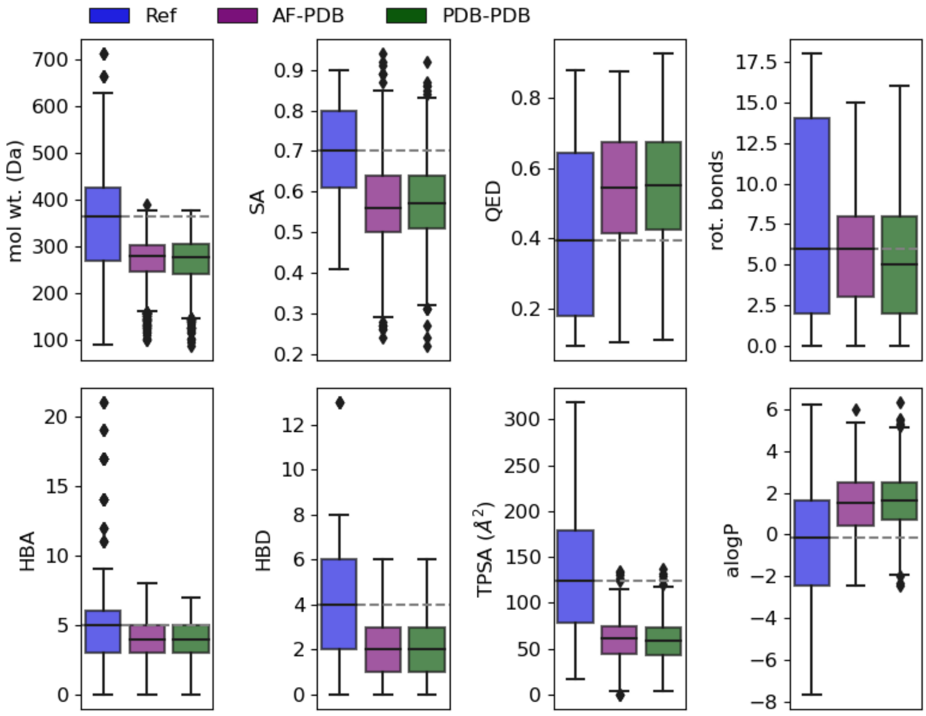


**Figure S5: Properties of ligands generated from AlphaFold vs. PDB receptors.** Distributions of properties for molecules generated for the AlphaFold test set receptors using the AF predicted structure (AF) and the PDB-derived crystal structure (PDB). For comparison, property distributions also shown for the crystal ligands (Ref) along with medians of crystal ligand distributions (grey dashed line). (*mol. wt.: molecular weight, SA: synthetic accessibility, QED: drug-likeness, rot. bonds: rotatable bonds, HBD: H-bond donors, HBA: H-bond acceptors, TPSA: topological polar surface area, alogP: hydrophobicity*)

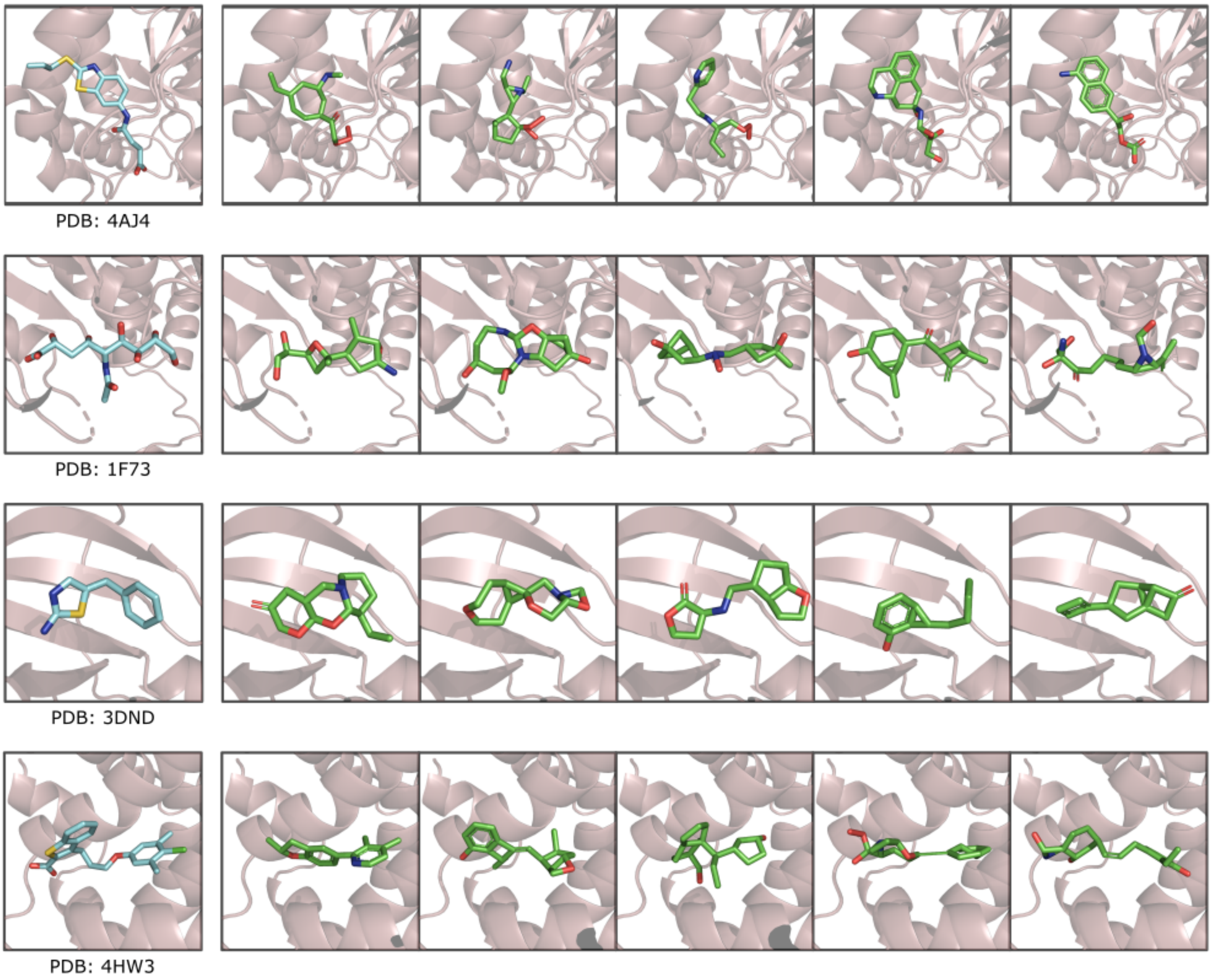


**Figure S6: Random generated molecules from prior sampling.** Random sets of generated molecules targeted at PDB crystal receptors sampled from the DrugHIVE prior.

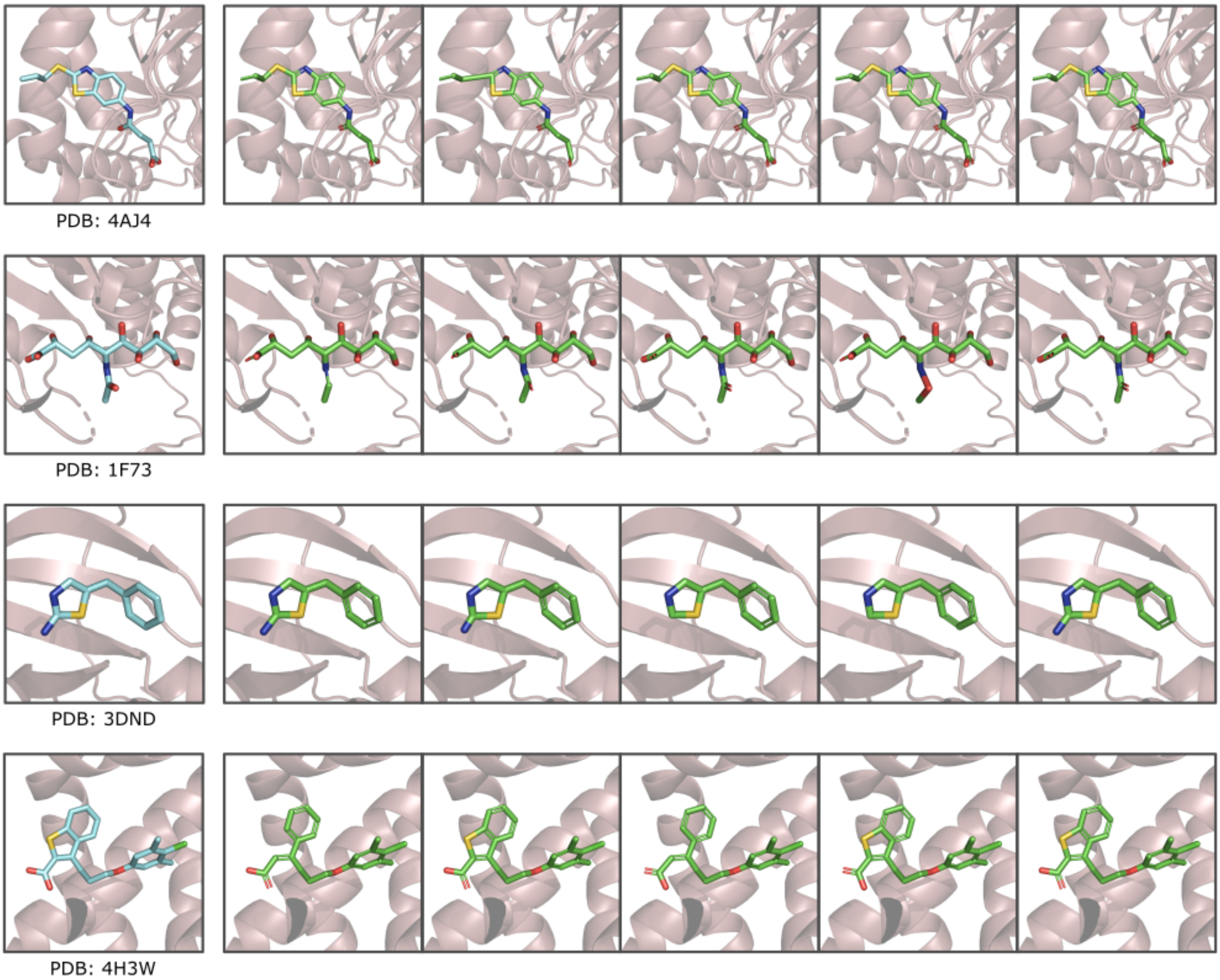


**Figure S7: Random generated molecules from posterior sampling.** Random sets of generated molecules targeted at PDB crystal receptors sampled from the DrugHIVE posterior ($\beta$=0) for the crystal ligand.

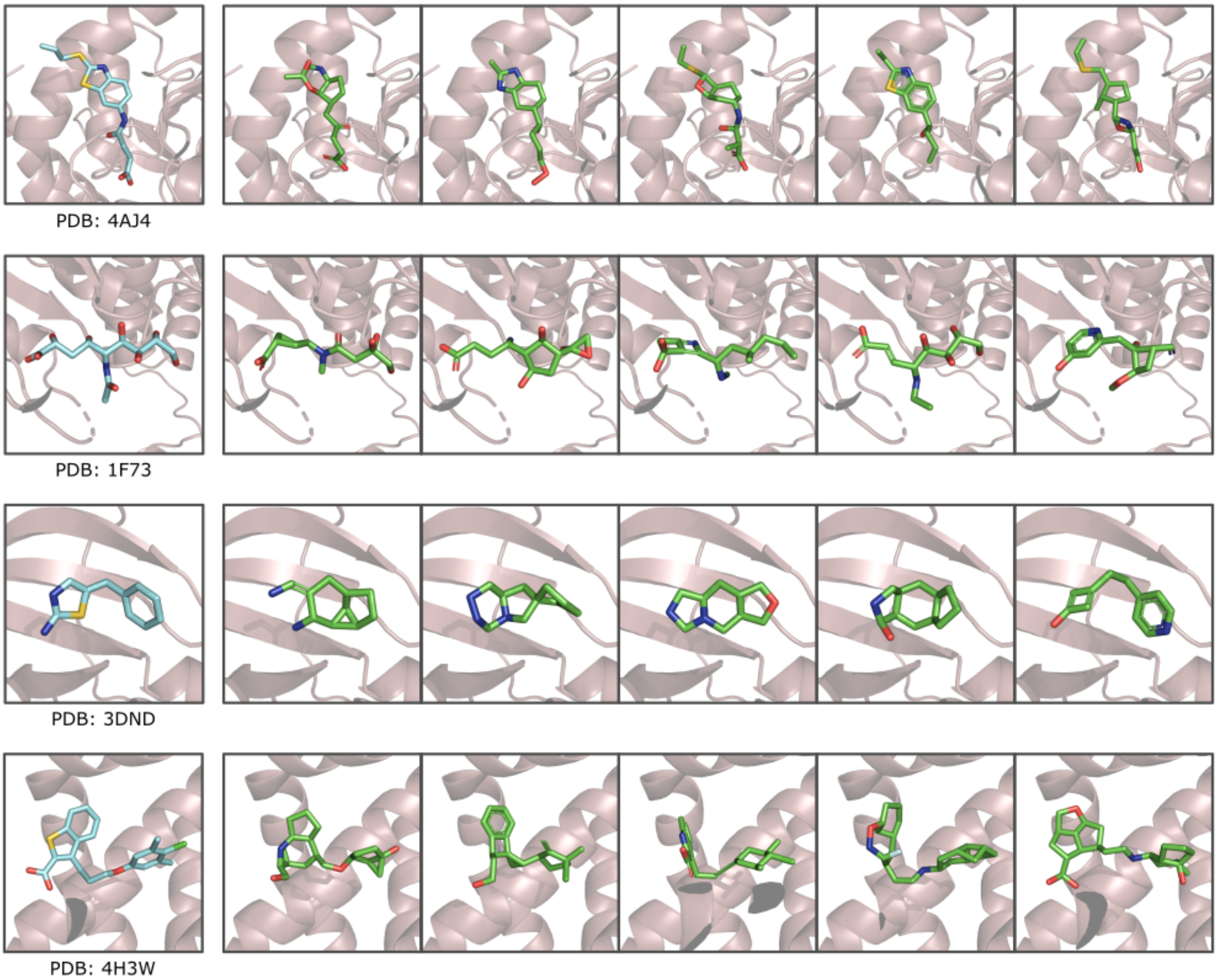


**Figure S8: Random generated molecules from prior-posterior sampling ($\beta$=0.5).** Random sets of generated molecules targeted at PDB crystal receptors sampled from DrugHIVE using prior-posterior sampling with interpolation factor $\beta$=0.5, starting from the crystal ligand structure.

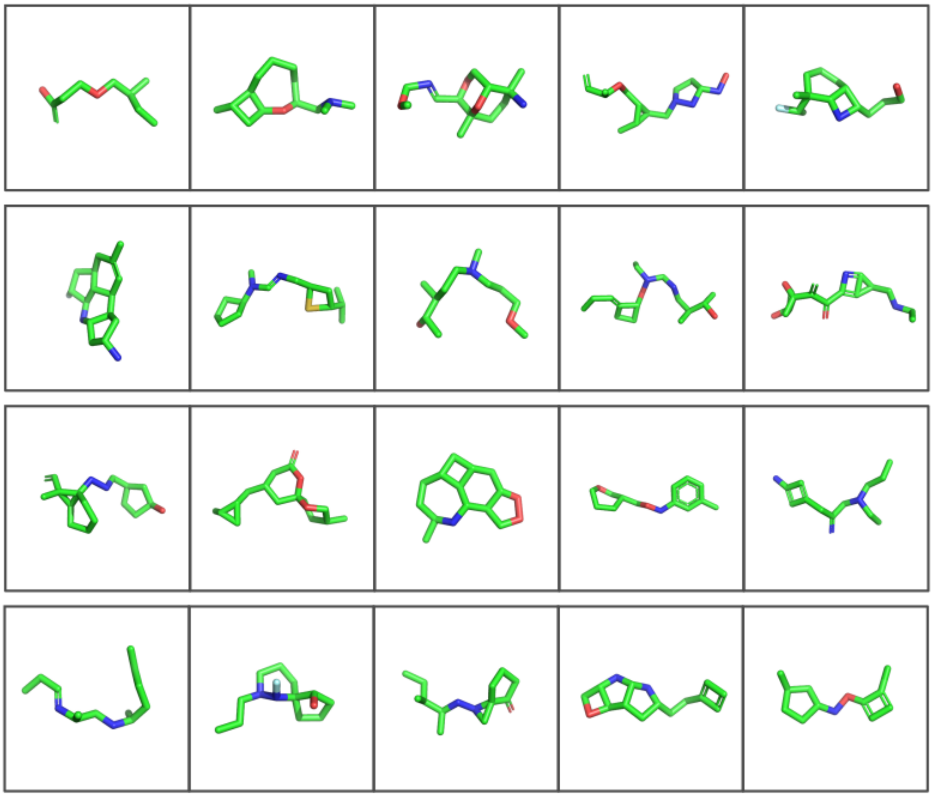

**Figure S9: Random generated molecules from prior sampling without receptors.** A random set of generated molecules with no receptor sampled from the DrugHIVE prior.
